# Genetic signatures in the highly virulent subtype B HIV-1 conferring immune escape to V1/V2 and V3 broadly neutralizing antibodies

**DOI:** 10.1101/2024.03.13.584899

**Authors:** Manukumar Marichannegowda, Alonso Heredia, Yin Wang, Hongshuo Song

## Abstract

HIV-1 is considered to become less susceptible to existing neutralizing antibodies over time. Our study on the virulent B (VB) HIV-1 identified genetic signatures responsible for immune escape from broadly neutralizing antibodies (bNAbs) targeting V1/V2 and V3 glycan epitopes. We found that the absence of N295 and N332 glycans in the high mannose patch, which are crucial for neutralization by V3 glycan bNAbs and are typically conserved in subtype B HIV-1, is a notable feature in more than half of the VB variants. Neutralization assays confirmed that the loss of these two glycans in VB HIV-1 leads to escape from V3 glycan bNAbs. Additionally, all VB variants we investigated have an insertion in V2, contributing to immune escape from V1/V2 bNAbs PG9 and PG16. These findings suggest potential co-evolution of HIV-1 virulence and antigenicity, underscoring the need to monitor both the pathogenicity and neutralization susceptibility of newly emerged HIV-1 strains.

## Introduction

The high mutation rate, high recombination rate, and rapid turnover of HIV-1 in an infected individual allow the virus to explore the sequence space extensively under different mechanisms of host defense. Consequently, extraordinary genetic diversity of HIV-1 is formed over more than 40 years of pandemic. Currently, a total of 9 subtypes and 157 circulating recombination forms (CRFs) of the group M HIV-1 are circulating across the globe^1^. The genetic diversity of HIV-1 is continuously evolving over time, which represents a major challenge in HIV-1 vaccine development.

While the extensive genetic diversity of HIV-1 has become apparent early during the pandemic, how such genetic diversity translates into virus phenotypic traits, and in turn influence the disease outcome remains relatively poorly understood. Emerging evidence showed that even within a single HIV-1 clade, new genetic cluster(s) with enhanced virulence could emerge. One example is the identification of certain CRF01_AE HIV-1 clusters with high prevalence of CXCR4 variants, which correlates with low CD4^+^ T cell count and poor immune reconstitution on ART^2,3^. Another example is the identification of the highly virulent subtype B (VB) HIV-1 in Netherlands in 2022^4^. The VB variants, initially identified in 109 participants in the BEEHIVE and ATHENA projects, led to 3.5 - 5.5 folds increase in the set point viral load (VL), and a 2-fold increased rate of CD4 decline compared to other subtype B viruses^4^. Sequence analysis suggests that the VB HIV-1 originated around 1998 through *de novo* mutations rather than recombination^4^. Transmission cluster study indicates potentially increased transmissibility of the VB strains^4^. Of note, following the initial detection of the VB variants in Netherlands, three VB cases of Polish origin and one case of Belgium origin were identified in 2023^5^, which raised the concern of the spread of the VB HIV-1 to other regions of the world.

HIV-1 is considered to adapt to both the cellular and humoral immunity over time at a population level^6,7^. Particularly, decreased susceptibility to existing bNAbs targeting different envelope (*env*) domains was reported for subtype B, subtype C, and CRF01_AG HIV-1^8–13^. Because bNAbs are currently under intensive study as preventative, therapeutic, as well as functional cure strategies, it is necessary to evaluate the susceptibility of newly emerged HIV-1 variants to currently available bNAbs and to determine potential mechanisms responsible for immune evasion. Among the total of 250 amino acid substitutions in the VB HIV-1 as compared to the Dutch subtype B consensus sequence, 30 were predicted to associate with escape to cytotoxic T cells (CTL)^4^. However, whether the VB HIV-1 are associated with immune escape to bNAbs has not been addressed so far. In the current study, we identified genetic signatures in the V2 and V3 loops of the VB variants associated with resistance to neutralization, and functionally demonstrated that these genetic signatures are responsible for immune escape to V1/V2 and V3 glycan bNAbs.

## Results

### The N295 glycan site and the N332 glycan supersite are not conserved in the VB HIV-1

The N295 glycan site and the N332 glycan “supersite” in the V3 high mannose patch are important determinants for neutralization susceptibility to V3 glycan bNAbs^14–19^. When analyzing the *env* sequences of the 17 VB strains, we observed high frequency of amino acid substations at the N295 and N332 positions, which led to the loss of the conserved N295 and N332 glycans (Figure 1A). Of the 17 VB variants, 12 variants (70.6%) lacked the N295 glycan, mostly due to amino acid substitutions at the N295 position (Figure 1A). Nine of the 17 VB variants (52.9%) lacked the N332 glycan supersite. Of note, in all these nine VB variants, the N332 glycan shifted to the N334 position (Figure 1A), a previously identified mechanism for immune escape to V3 glycan bNAbs^18,19^. Six VB variants (35.3%) lost both the N295 and N332 glycans, and only two VB variants (11.8%) contained both glycans (Figure 1A). In addition to the N295 and N332 glycans, the D325N substitution in the V3^324^ GDIR^327^ motif, a mutation associated with immune escape to several V3 glycan bNAbs^19–21^, was found in 9 VB variants (52.9%) which lacked the N295 and/or N332 glycans (Figure 1A). Phylogenetic analysis showed that the two VB variants containing intact N295 and N332 glycans, and the three variants containing the N295 glycan but lacking the N332 glycan, were closely related in the tree, respectively, indicating potential transmission relationship of these variants (Figure S1).

**Figure 1.**
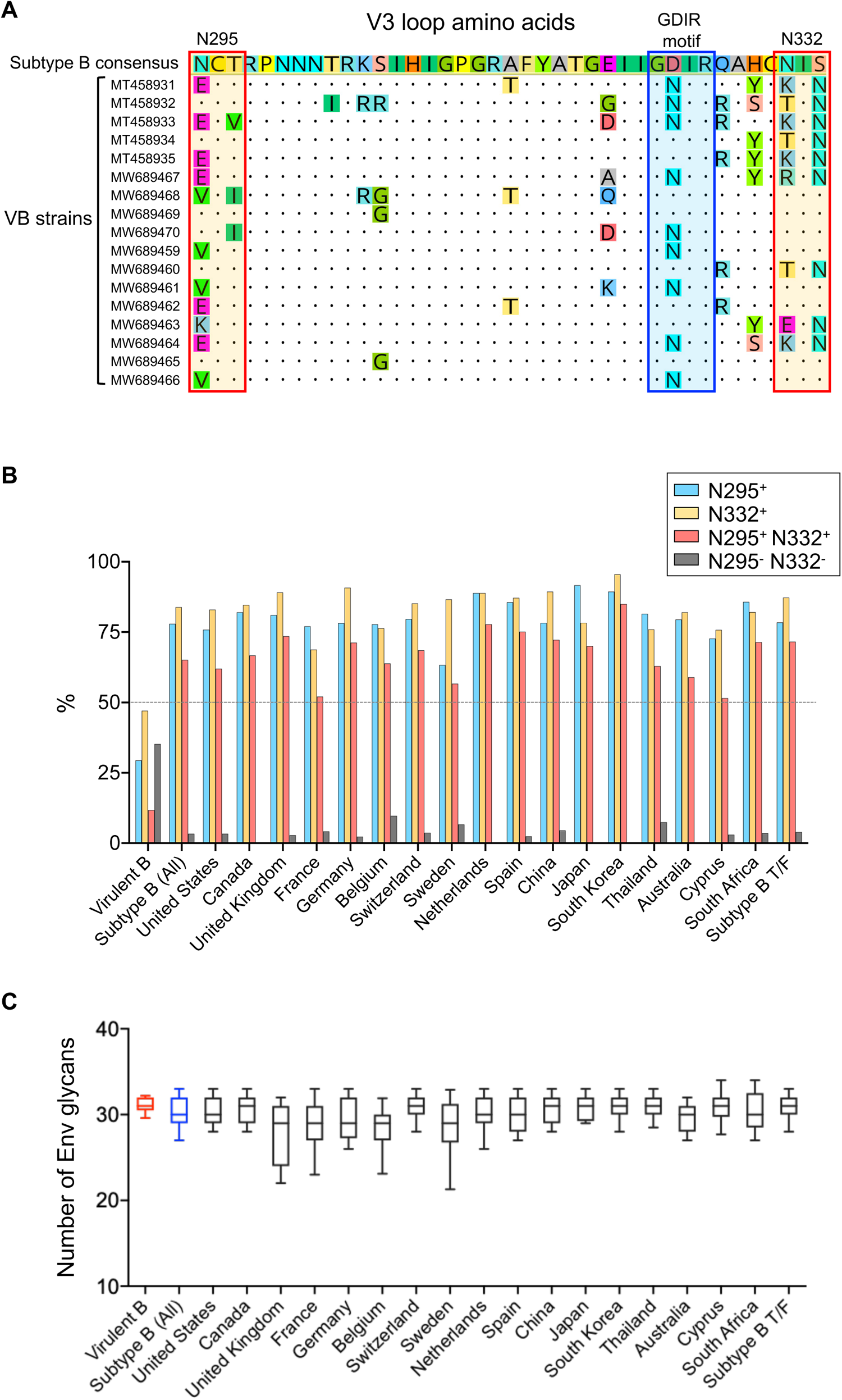
The N295 and N332 glycan sites in the V3 high mannose patch are not conserved in the VB HIV-1. (A) V3 amino acid alignment of 17 publicly available VB sequences identified in the BEEHIVE cohort. The GenBank accession number is used as the name of each sequence. The subtype B consensus sequence is used as the reference sequence. The N295 and N332 glycan sites are indicated by red boxes. The GDIR motif is indicated by blue box. (B) Frequency of the N295 and N332 glycan sites in the VB variants compared to other subtype B viruses from 17 different countries. A total of 102 subtype B T/F viral sequences were analyzed separately. (C) Comparison of the total number of N-linked glycans in envelope between the VB variants and other subtype B viruses. The data of the VB variants and the combined data of all other subtype B viruses are shown in red and blue, respectively. For each group, the line shows the median, and the box and whisker indicate the 10^th^ and 90^th^ percentiles and the range, respectively.

To determine whether the low frequency of the N295 and N332 glycan sites is unique to the VB variants, we analyzed the prevalence of these two glycans in other subtype B viruses identified in different countries. A total of 17 countries, which had at least 20 subtype B *env* sequences available in the Los Alamos HIV Sequence Database, were included in the analysis (Figure 1B). We also included 102 subtype B transmitted/founder (T/F) viruses identified in a previous study^22^. The analysis showed that both the N295 and N332 glycan sites were conserved in other subtype B viruses from different countries (Figure 1B). Among the total of 3,240 subtype B HIV-1 sequences analyzed, 78.0% contained intact N295 glycan site (ranging from 63.3% to 91.7% when the data was stratified by countries), and 83.9% contained intact N332 glycan site (ranging from 68.8% to 95.6%). Only 3.3% of the 3,240 subtype B HIV-1 sequences lacked both the N295 and N332 glycans (ranging from 2.3% to 9.7%), while 65.1% of them contained both glycans (ranging from 51.5% to 85.0%) (Figure 1B). These two glycan sites were similarly conserved in the 102 subtype B T/F viruses (Figure 1B).

We next sought to determine whether the VB HIV-1 had an overall low number of N-linked glycan sites in the envelope protein. The VB variants contained an average of 31 N-linked glycan sites in their envelope protein, while an average of 30 N-linked glycan sites were present in other subtype B viruses (Figure 1C). Therefore, the low frequency of the N295 and N332 glycans in the VB variants was not due to an overall low number of N-linked glycan sites in the envelope. Indeed, another 13 N-linked glycans in gp120, which are conserved (with > 50% frequency) in subtype B HIV-1 as reported previously^23^, remained conserved in the VB strains (Figure S2). Taken together, the low frequency of the N295 and N332 glycan sites was uniquely observed for the VB HIV-1, which was not due to an overall low number of N-linked glycan sites in the VB envelope.

### The VB variants have relatively shorter V1 region and contain an insertion in V2 region

The unusually low frequency of the N295 and N332 glycan sites, as well as the high frequency of the D235N substitution in the ^324^GDIR^327^ motif of the VB variants, raised the possibility that the VB HIV-1 could be associated with immune escape to neutralizing antibodies targeting the variable loops. Because the length of the *env* variable loops could also impact neutralization susceptibility, we compared the number of amino acids in each variable loop of the VB variants with other subtype B viruses. The VB variants had a relatively shorter V1 region (an average of 16 amino acids, ranging from 11 to 21) in comparison to other subtype B viruses (an average of 20 amino acids) (Figure 2A). Despite the difference in the length of V1, both the VB HIV-1 and other subtype B viruses contained an average of two N-linked glycans in the V1 loop (Figure S3). In comparison to other subtype B viruses, the VB variants had an obviously longer V2 loop (an average of 45 amino acids in comparison to an average of 40 amino acids in other subtype B viruses) (Figure 2B). Sequence alignment identified that all 17 VB strains contained an insertion in V2 at the same location (downstream of the HXB2 location 184) (Figure 2C). The number of amino acids in this insertion varied from 4 to 9 across different VB strains (Figure 2C). Of note, this insertion provided an additional N-linked glycan site to the V2 loop of the VB variants as compared to the subtype B consensus sequence (Figure S3). No significant difference in the length of the V3, V4 and V5 regions were observed between the VB variants and other subtype B viruses (Data not shown).

**Figure 2.**
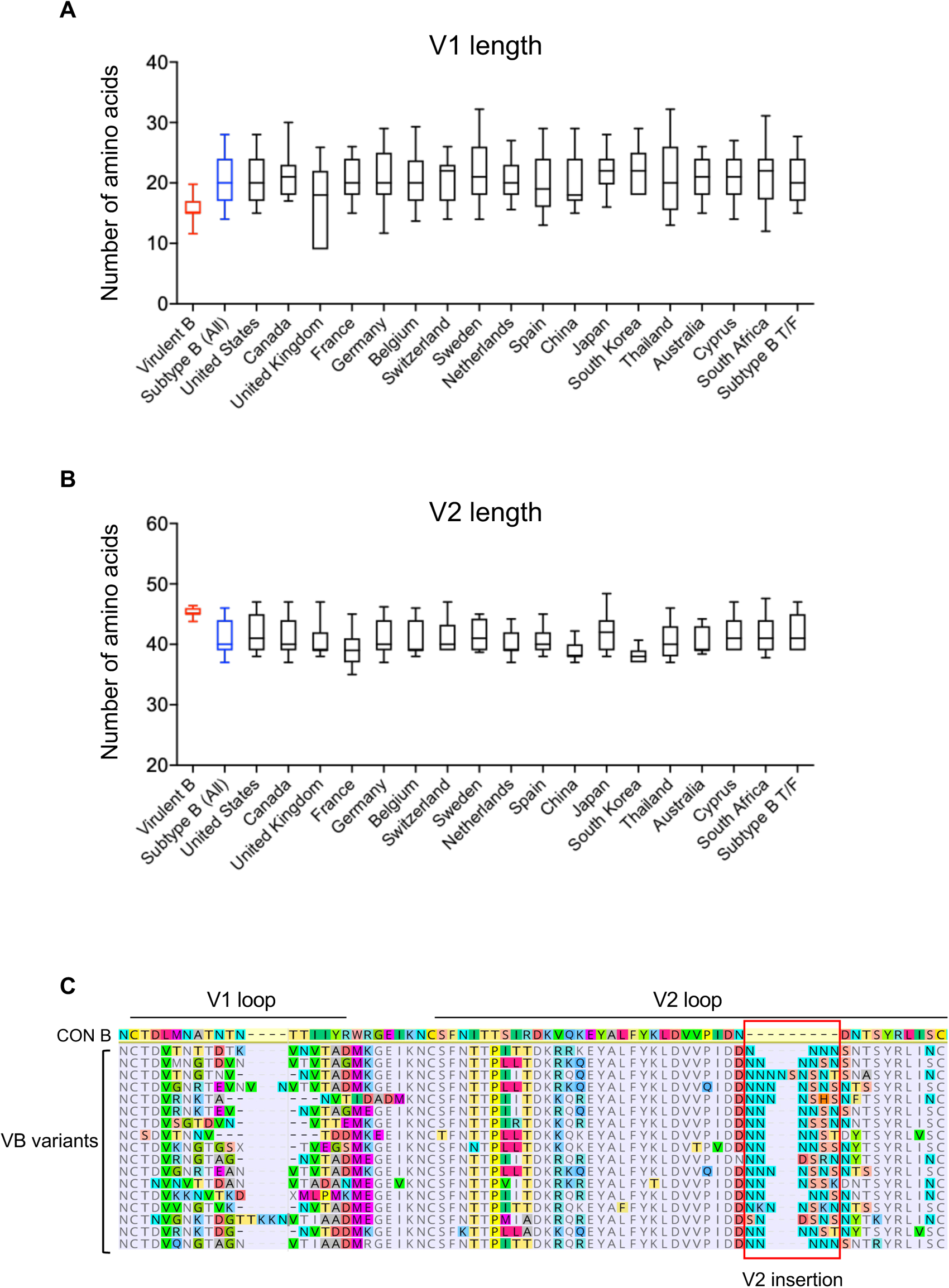
The VB variants have relatively shorter V1 loop and longer V2 loop due to an insertion in V2. (A) Comparison of the length of the V1 region between the VB variants and other subtype B viruses from different countries. (B) Comparison of the length of the V2 region between VB variants and other subtype B viruses from different countries. In both panel A and panel B, the line shows the median, while the box and whisker indicate the 10^th^ and 90^th^ percentiles and the range of each group, respectively. (C) Amino acid alignment of the V1/V2 loops of the VB variants. The subtype B consensus sequence (CON B) is used as the reference sequence. The insertion in the V2 region of the VB variants is indicated by the red box.

### The loss of the N295 and N332 glycan sites and the insertion in V2 confer immune escape to bNAbs

To functionally characterize whether the genetic signatures identified in the VB variants were responsible for immune escape to neutralizing antibodies, we determined the neutralization susceptibility of four representative VB strains (MT458931, MT458935, MW689460, and MW689465) to a panel of bNAbs targeting different HIV-1 *env* domains. The VB variants MT458931 and MT458935, which lacked both the N295 and N332 glycans, were completely resistant to the V3 glycan bNAbs PGT121, PGT126, PGT128, and 2G12, regardless of whether they carried the D325N mutation (Figure 3A-3B). Variant MW689460, which had the intact N295 glycan but lacked the N332 glycan, was also resistant to all tested V3 glycan bNAbs (Figure 3A-3B). Variant MW689465, which contained both the N295 and N332 glycans, were sensitive to all tested V3 glycan bNAbs (Figure 3A-3B).

**Figure 3.**
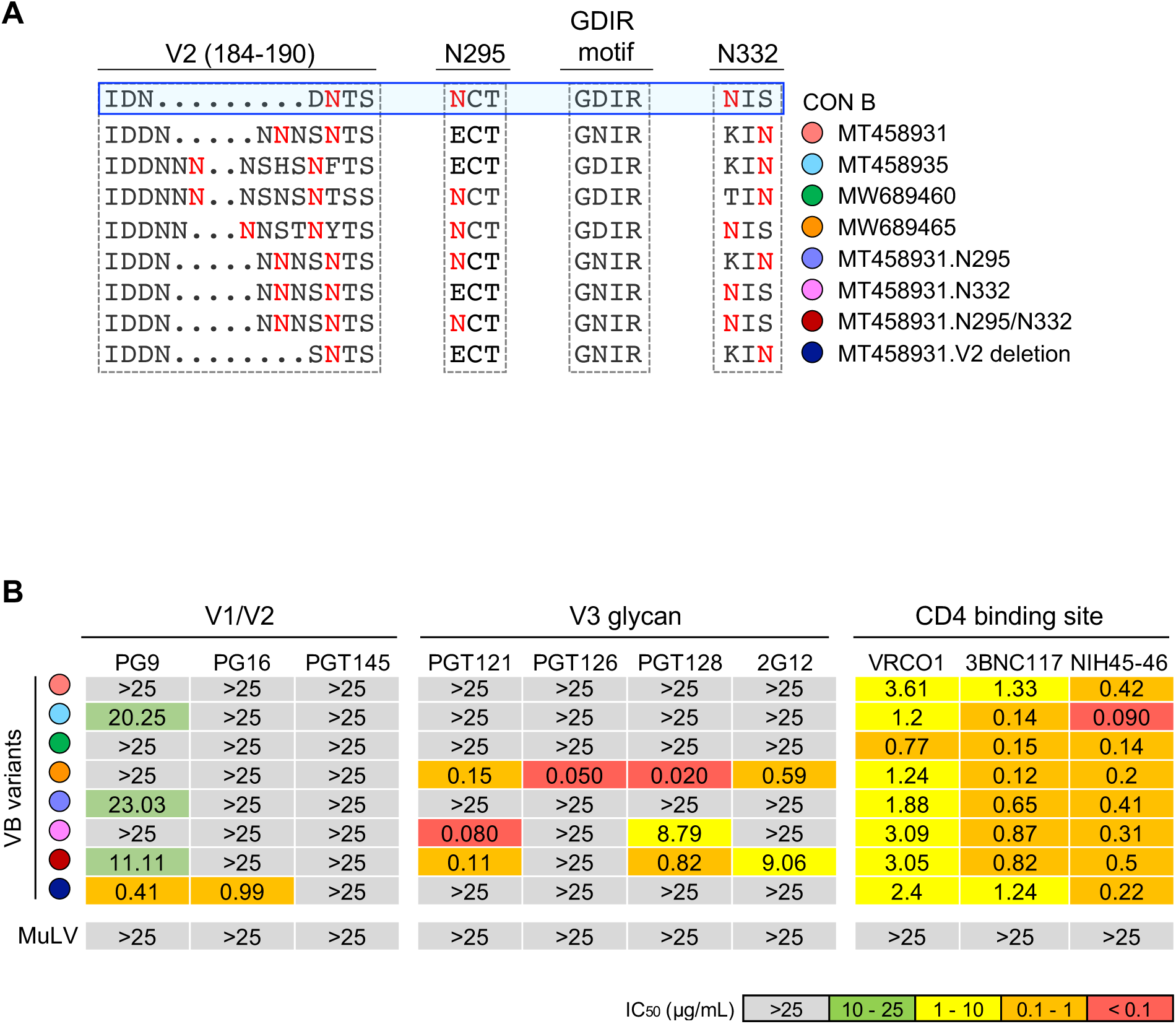
Impact of the N295 and N332 glycans and the insertion in V2 on neutralization susceptibility to broadly neutralizing antibodies (bNAbs) targeting different HIV-1 *env* domains. (A) Amino acid alignments of the V2 insertion stie, the N295 and N332 glycan sites, and the V3 GDIR motif of four representative VB strains and four mutants based on the variant MT458931 (Variant “MT458931.N295” has restored N295 glycan. Variant “MT458931.N332” has restored N332 glycan. Variant “MT458931.N295/N332” has both the N295 and N332 glycans restored. Variant “MT458931.V2 deletion” has three amino acids deletion in the V2 insertion). The subtype B consensus sequence (CON B) is used as the reference sequence. The N-linked glycan sites are highlighted in red. Different VB variants are color coded. (B) Neutralization susceptibility of different VB variants to a panel of bNAbs targeting different HIV-1 *env* domains. The number shows the IC_50_ (µg/mL) of each bNAb. The MuLV was used as negative control in the neutralization assay.

We further determined the contributions of the N295 and N332 glycans to neutralization susceptibility by “restoration of function” in the genetic background of MT458931, which lacked both glycans and was resistant to all V3 glycan bNAbs (Figure 3A-3B). Restoration of the N295 glycan alone could not restore the neutralization sensitivity to any of the V3 glycan bNAbs (Figure 3A-3B). Restoration of the N332 glycan alone restored the neutralization sensitive to PGT121 and PGT128 (Figure 3A-3B). Restoration of both the N295 and N332 glycans rendered the virus susceptible to 2G12 in addition to PGT121 and PGT128 (Figure 3A-3B). Additionally, the variant with both glycans had approximately 10-fold increased sensitivity to PGT128 as compared to the variant with the N332 glycan alone (Figure 3A-3B). These results indicated that the cooperation between the N295 and N332 glycans could play an important role in determining virus susceptibility to 2G12 and PGT128.

All four VB variants we investigated were resistant to the V1/V2 bNAbs PG9, PG16, and PGT145, regardless of the length of the V1 loop (Figure 3B). However, we found that the N156 and N160 glycans in the V2 loop, which are essential for neutralization by PG9, PG16, and PTG145^24–27^, were both conserved in the VB strains (Figure S2). This indicated that the VB variants might exploit other mechanism(s) to evade these V1/V2 bNAbs. Because an increased length of the variable loop, as well as insertion of N-linked glycans, are both potential mechanisms for neutralization escape^7^, we determined the impact of the V2 insertion on neutralization susceptibility. Three amino acids were deleted in the V2 insertion in variant MT458931, which removed the additional N-linked glycan in this insertion region (Figure 3A). The variant with this V2 deletion became sensitive to PG9 and PG16 (IC_50_ = 0.41 µg/mL and 0.99 µg/mL, respectively) (Figure 3B). However, the susceptibility to PGT145 could not be restored by this deletion. The deletion in V2 did not impact virus sensitivity to bNAbs targeting the V3 and the CD4 binding site (Figure 3B). The four VB variants we investigated showed similar level of neutralization sensitivity to the CD4 binding site bNAbs VRC01, 3BNC177, and NIH45-46 as compared to the contemporary subtype B viruses reported in a recent study^12^.

### The VB variants rely on CCR5 to enter primary CD4^+^ T cells and are sensitive to the CCR5 inhibitor Maraviroc

All except one VB variants were predicted to be CCR5 tropic based on the V3 sequences in the previous study^4^. However, their coreceptor usage has not been functionally determined. We investigated the coreceptor usage of the four VB variants in a panel of NP-2 cell lines expressing CCR5, CXCR4, as well as another seven alternative coreceptors which could be used by HIV-1 (CCR1, CCR2b, CCR3, CCR6, CCR8, APJ, GPR15). All four VB strains used CCR5 with similar efficiency in NP-2 CCR5 cell line (Figure 4A). Although previous study showed that the GDIR motif comprises part the CCR5 binding region, and mutations in this motif could compromise CCR5 binding^21^, we did not observe a compromised CCR5 usage for the variant (MT458931) which carried the D235N mutation in the GDIR motif (Figure 4A). Different from the control virus CH040 (a subtype B T/F virus), three of the four VB variants (except for MT458931) could use CCR3 with high efficiency in NP-2 CCR3 cell line (Figure 4A). Restoration of the N295 and/or N332 glycans did not have observable impact on coreceptor usage (Figure 4A). All four VB variants were sensitive to the CCR5 inhibitor Maraviroc in NP-2 CCR5 cell line (Figure S4). The average IC_50_ of Maraviroc, as determined in NP-2 CCR5 cell line, was 11.9 nM for the VB strains, which was approximately 2-fold higher than the subtype B T/F virus CH040 (5.6 nM) (Figure S4).

**Figure 4.**
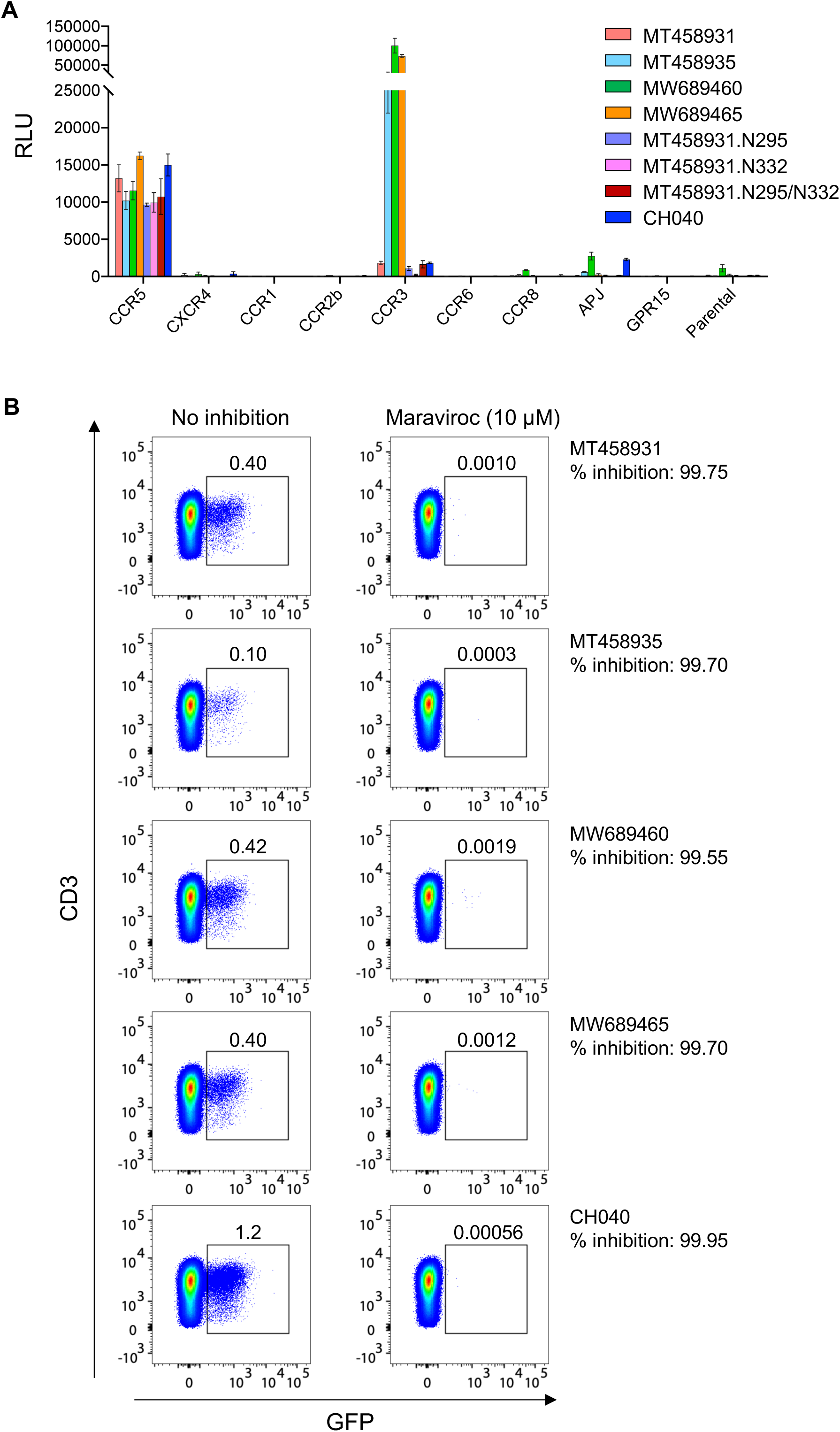
Coreceptor usage of the VB variants in NP-2 cell lines and sensitivity to Maraviroc in primary CD4^+^ T cells. (A) Coreceptor usage of four representative VB variants in a panel of NP-2 cell lines expressing different HIV-1 coreceptors. The potential impact of the N295 and N332 glycans on coreceptors usage was determined using three mutants based on the VB strain MT458931, as described in Figure 3. The subtype B T/F virus CH040 was used as control. All experiments were performed in triplicate. The error bar shows the standard deviation (SD). (B) Infectivity of four VB variants in primary CD4^+^ T cells and the sensitivity to Maraviroc inhibition. Stimulated CD4^+^ T cells with and without Maraviroc (10 µM) pre-treatment were infected by each VB pseudovirus containing the GFP reporter in the backbone. The percentage of GFP positive cells were determined by flow cytometry three days after infection. The subtype B T/F virus CH040 was used as control. The percentage of Maraviroc inhibition was determined by comparing the percentage of GFP positive cells between infections with and without Maraviroc inhibition.

The observation that certain VB variants exhibited high efficiency in using CCR3 *in vitro* raised the question of whether they could enter primary CD4^+^ T cells through alternative coreceptor(s). To determine this, we infected primary CD4^+^ T cells with VB pseudoviruses containing the GFP reporter and assessed their sensitivity to Maraviroc inhibition. In the absence of Maraviroc, between 0.10% to 0.42% of primary CD4^+^ T cells were GFP positive after three days of infection (Figure 4B). When primary CD4^+^ T cells were pre-treated with 10 µM of Maraviroc, over 99.5% of viral infectivity was inhibited for all four VB strains (Figure 4B). For the subtype B T/F virus CH040, the same concentration (10 µM) of Maraviroc could inhibit 99.95% of viral infectivity (Figure 4B). These results demonstrate that the VB variants rely on CCR5 for infecting primary CD4^+^ T cells. However, whether the extremely low level of residual infectivity was mediated by alternative coreceptor(s), and whether it is biologically relevant *in vivo* remain to be determined.

## Discussion

Over the course of the pandemic, HIV-1 of different subtypes have been shown to become less susceptible to existing bNAbs targeting nearly all *env* domains^8–13^. In the current study, we identified clear genetic signatures in the recently emerged VB HIV-1 that are responsible for immune escape to bNAbs targeting the V1/V2 and V3 glycan epitopes. The evolutionary trend of HIV-1 towards resistance to currently available bNAbs, as well as the potential co-evolution of HIV-1 antigenicity and virulence, should be better investigated in the future in order to inform the development of preventative and therapeutic approaches based on bNAbs.

The VB variants have evolved multiple immune escape pathways, which are likely driven by neutralizing antibodies targeting different epitopes across the V1/V2 and V3 loops. The escape mechanisms in the V3 high mannose patch includes the shift of the N332 glycan supersite to the N334 position, the loss of the N295 glycan, as well as the substitutions in the GDIR motif. Neutralization assays demonstrated that the N332 glycan supersite is likely a key determinant for neutralization sensitivity to V3-glycan bNAbs in the genetic context of the VB strains, while the cooperation between the N295 and N332 glycans are required for neutralization susceptibility to 2G12. Despite the conservation of the N156 and N160 glycans in the V2 loop, the VB variants remained resistant to V1/V2 bNAbs PG9, PG16, and PGT145. Deletion of the V2 insertion, which exists in all 17 VB variants and carries an additional N-linked glycan, was sufficient to restore the neutralization sensitivity to V1/V2 bNAbs PG9 and PG16. These findings together provide evidence that during the evolutionary history of the VB variants, the viruses were likely experienced strong selection pressures in the infected individuals exerted by neutralizing antibodies with similar specificities to the V1/V2 and V3 bNAbs. It is possible that the escape phenotype of the VB variants is a consequence of “founder effect”. That is, the ancestor of the currently circulating VB variants already carried these escape mutations. Over time, reversions might occur in the absence of the selection, which led to the polymorphisms currently observed at the N295 and N332 positions. Another possibility is that the VB strains, after originated, have been under strong humoral immune pressures targeting the V1/V2 and V3 loops in the infected individuals, which led to a higher than usual frequency of immune escape mutations. If this scenario is true, it would be interesting to determine whether the ancestral VB strain has unique antigenicity feature that could elicit stronger V1/V2 and V3 humoral responses than other subtype B viruses.

Another interesting finding is that three of the four VB variants we investigated exhibited high efficiency in using CCR3 *in vitro*. Although CCR3 does not contribute to efficient viral entry into the primary CD4^+^ T cells, as demonstrated by Maraviroc inhibition assay, its potential role in viral pathogenicity in other cell types deserves further investigation. Particularly, CCR3 is known to contribute to efficient HIV-1 infection of the microglia^28^, the major target cells of HIV-1 in the central nervous system (CNS)^29,30^. HIV-1 isolated from the CNS, such as the YU2 strain, exhibits high efficiency in using CCR3^31^. Therefore, it would be necessary to determine in the future whether the VB variants could infect the microglia with high efficiency, thereby having an enhanced neurovirulence.

While the enhanced CD4 virulence of the VB HIV-1 was considered as a “unfamiliar” mechanism associated with CCR5 tropism^4^, we hypothesize that the possibility that the VB variants could undergo early coreceptor switch in the infected individuals may also need to be explored. Our recent study, by tracking coreceptor switch in two participants in the RV217 acute infection cohort, provides evidence that coreceptor switch could be driven by autologous NAbs^32^. We hypothesized that coreceptor switch could function as a compensatory mechanism to maintain viral entry capacity, when the CCR5 usage is impaired by immune escape mutations driven by autologous NAbs^32^. Of note, both participants who underwent early coreceptor switch were infected by T/F viruses with unique glycan arrangement at the N295 and N332 positions (one T/F virus did not have the N332 or N334 glycan, and another T/F virus did not have the N295 glycan). It is possible that the absence of the N295 and/or N332 glycans in the transmitted virus could expose the V3 loop to early humoral immune pressures, thereby increasing the chance of coreceptor switch. It is important to point out that, in all participants who underwent coreceptor switch in the RV217 cohort, the X4 variants existed at low frequency in plasma compared to the co-existing R5 viruses, which is likely due to their altered CD4 subset tropism towards the naïve and central memory CD4 subsets^32^. Because the sequences of the 17 VB variants are likely representing the predominant viral population in the plasma, the existence of low-frequency, CXCR4 variants in VB HIV-1 infected people could not be completely excluded. Future studies by deep sequencing the plasma viral quasispecies, or by sequencing the viruses in the central memory and naïve CD4 subsets, could provide important information on whether coreceptor switch have occurred in people infected by the VB HIV-1.

The limitation of the current study is that we only analyzed the 17 VB variants originally identified in the BEEHIVE cohort^4^. While more cases of VB HIV-1 infections were identified previously^4^, the viral envelope sequences are not publicly available at this time. The availability of more VB *env* sequences in the future will be useful to comprehensively determine the prevalence of neuralization resistant variants in the VB HIV-1.

## Acknowledgments

This study was supported by Institute of Human Virology at the University of Maryland School of Medicine, and National Institutes of Health (NIH) grants R21AI147893 and R01AI120008.

## Methods

### HIV-1 sequence data

The 17 virulent subtype B (VB) HIV-1 sequences reported previously in the BEEHIVE project^4^ were downloaded from the Los Alamos HIV Sequence Database. In addition, a total of 3,240 subtype B HIV-1 *env* sequences from 17 countries were retrieved from the Los Alamos HIV Sequence Database (we excluded the countries which had less than 20 subtype B *env* sequences available in the database). We also included 102 HIV-1 subtype B transmitted/founder *env* sequences reported in a previous study^22^.

### Genetic analysis

HIV-1 *env* sequences were aligned by the Gene Cutter tool in the Los Alamos HIV Sequence Database (https://www.hiv.lanl.gov/content/sequence/GENE_CUTTER/cutter.html), followed by manual adjustment to obtain the optimal alignment. Sequence of each *env* variable loop was extracted using the Gene Cutter. N-linked glycosylation sites were analyzed by the N-GlycoSite tool in the Los Alamos HIV Sequence Database^33^. Sequence alignment was visualized using the Geneious software (https://www.geneious.com). Sequences with ambiguous codons and/or frameshift mutations in the variable loops were excluded from the analysis. Phylogenetic tree was constructed by MEGA6 using the maximum likelihood method^34^.

### Generation of HIV-1 envelope clones

The full-length *env* sequences of four representative VB strains MT458931, MT458935, MW689460, and MW68945 were chemically synthesized (GenScript, USA) and cloned into the expression vector pcDNA3.3-TOPO (Invitrogen, USA). Desired mutations were introduced by site-directed mutagenesis. All *env* clones were confirmed by sequencing.

### Pseudovirus preparation and titration

The pseudovirus stocks were prepared as previously described^35^. In brief, 2 μg of HIV-1 *env* clone was co-transfected with 4 μg of the pNL4.3-ΔEnv-vpr+-luc+ backbone (for coreceptor usage assay in NP-2 cell lines)^35^, or 4 μg of the pNL4.3-ΔEnv EGFP backbone^36^ (for viral entry assay in primary CD4^+^ T cells) into 293T cells using the FuGENE6 transfection reagent (Promega, USA). The cells were cultured at 37°C for 6 hours before the medium was completely replaced with fresh medium. The culture supernatants containing the pseudoviruses were harvested at 72 hours post transfection, aliquoted and stored at −80°C until use. The infectious titers (TCID_50_) of the pseudovirus stocks were determined in TZM-bl cell line.

### Neutralization assay

The neutralization activity of monoclonal antibodies (mAbs) was measured by using a luciferase reporter system in TZM-bl cells as previously described^37^. The mAbs were diluted at a 1:3 serial dilution from a starting concentration of 25 μg/mL. The virus stocks were diluted to a concentration that achieved approximately 150,000 relative luminescence units (RLU) in the TZM-bl cells (or at least 10 times above the background RLU of the cells control). The serial diluted mAbs were incubated with the diluted virus stocks for 1 hour at 37°C in duplicate before the TZM-bl cells were added. The IC_50_ (µg/mL) was determined as the mAb concentration at which the RLUs were reduced by 50% in comparison to the RLUs in the virus control wells after subtraction of the background RLUs in cell control wells.

### Coreceptor usage assay

The coreceptor usage of each virus was determined using a panel of NP-2 cell lines expressing CD4 together with different G protein-coupled receptors (CCR5, CXCR4, APJ, CCR3, CCR8, CCR1 and CCR2b) as previously described^35^. The parental NP-2 cell line expressing CD4 alone was used as a control. NP-2 cell lines expressing different coreceptors were seeded into a 96-well plate one day before infection at a density of 1 × 10^5^ cells per well. The next day, the cells were infected with approximately 200 TCID_50_ of each pseudovirus (MOI = 0.002) containing the pNL4.3-ΔEnv-vpr+-luc+ backbone. After 6 hours of infection at 37°C, the cells were washed three times with culture medium and cultured at 37°C for three days. At 72 hours post infection, the cells were lysed, and the infectivity was determined by measuring the RLUs in the cell lysates using the Britelite plus system (PerkinElmer, USA). Viral infection was considered positive if the RLU value was more than 5-fold higher than the background RLU value in parental NP-2 cell line. All experiments were performed in triplicate.

### Maraviroc inhibition assay in NP-2 CCR5 cell line

NP-2 CCR5 cells were seeded in a 96-well plate at a density of 1 × 10^5^ cells per well. The next day, the cells were pre-treated with 1:2 serial diluted Maraviroc (starting from 0.5 µM) at 37°C for 1 hour. The Maraviroc pre-treated cells were then infected with approximately 500 TICD_50_ of each pseudovirus containing the pNL4.3-ΔEnv-vpr+-luc+ backbone and cultured at 37°C for three days. The infected cells were lysed at day 3 after infection. The infectivity in each well was determined by measuring the RLU in the cell lysate using the Britelite plus system (PerkinElmer). The percentage of Maraviroc inhibition was determined by comparing the RLU values with the positive control wells without Maraviroc inhibition. The IC_50_ of Maraviroc was determined using a linear regression model.

### Determination of virus infectivity and sensitivity to Maraviroc in primary CD4^+^ T cells

Primary CD4^+^ T cells were purified from PBMCs of a healthy donor by negative selection (Miltenyi Biotec, USA). Purified CD4^+^ T cells were stimulated with 1 µg/mL soluble anti-CD3 (clone OKT3, eBioscience, USA) and 1 µg/mL soluble anti-CD28 (clone CD28.2, eBioscience, USA) in the presence of 50 IU/mL IL-2 (PeproTech, USA) for three days. One million of stimulated CD4^+^ T cells were infected by each pseudovirus (MOI = 0.1) containing the pNL4.3-ΔEnv EGFP backbone at 37°C for 4 hours. The infected cells were washed three times with RPMI1640 and were cultured in a 24-well plate in RPMI1640 containing 10% FBS and 50 IU/mL IL-2. Three days after infection, the percentage of GFP positive cells were determined by flow cytometry. To determine virus sensitivity to Maraviroc, one million stimulated CD4^+^ T cells were pre-treated with 10 µM Maraviroc at 37°C for 1 hour before infection with each pseudovirus as described above. The percentage of GFP positive cells were determined by flow cytometry three days after infection. The percentage of Maraviroc inhibition was determined by comparing the percentage of GFP positive cells between infections with and without Maraviroc inhibition.

## Supplemental information

**Figure S1.**
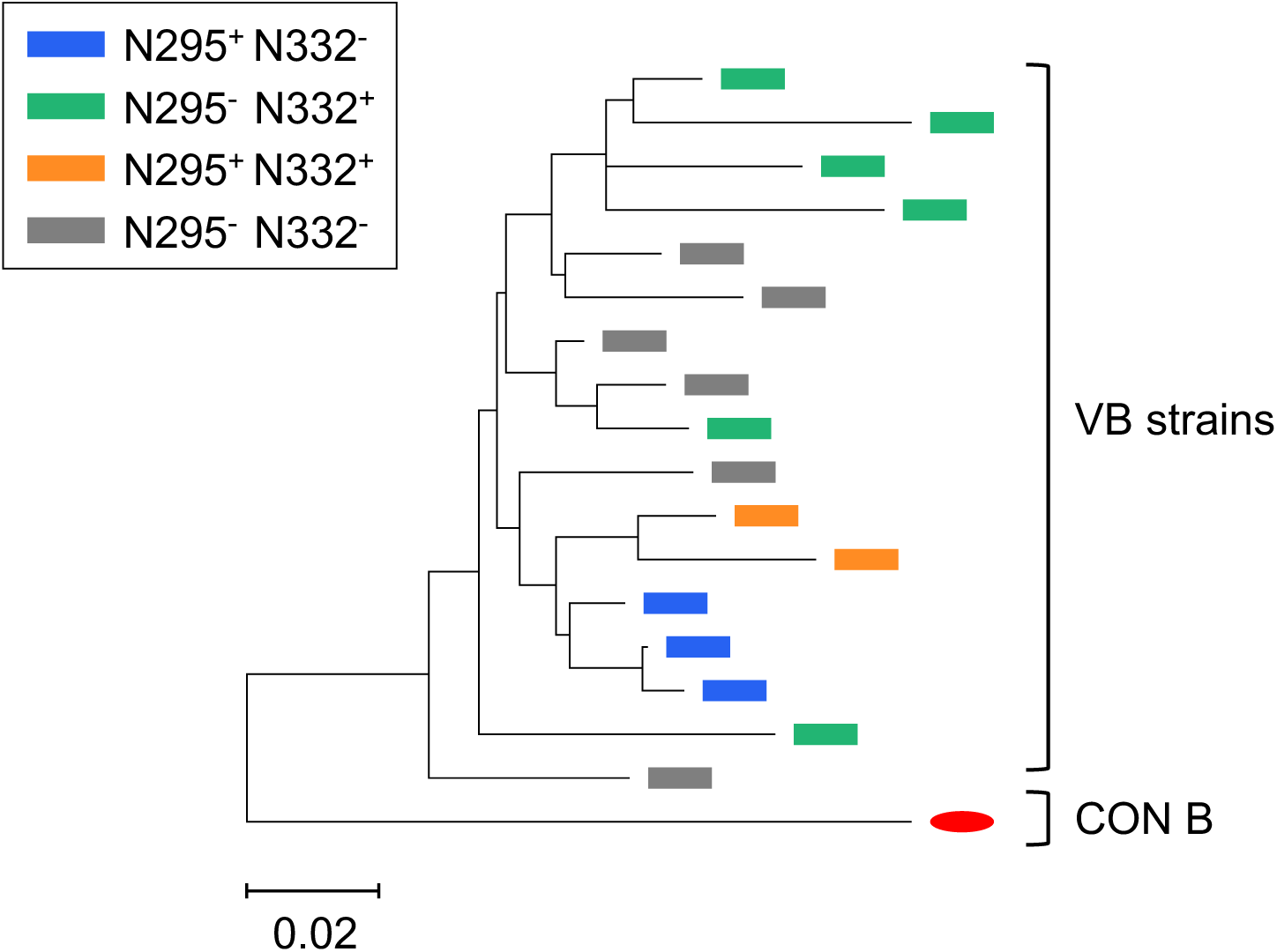
Phylogenetic relationship of the 17 VB HIV-1 variants. The *env* sequences of the 17 VB variants from the BEEHIVE project were used for phylogenetic analysis. The 17 VB variants are color coded based on the glycan status at the N295 and N332 positions. The subtype B consensus sequence (CON B) is shown in red. The tree was constructed by MEGA6 using the maximum likelihood model.

**Figure S2.**
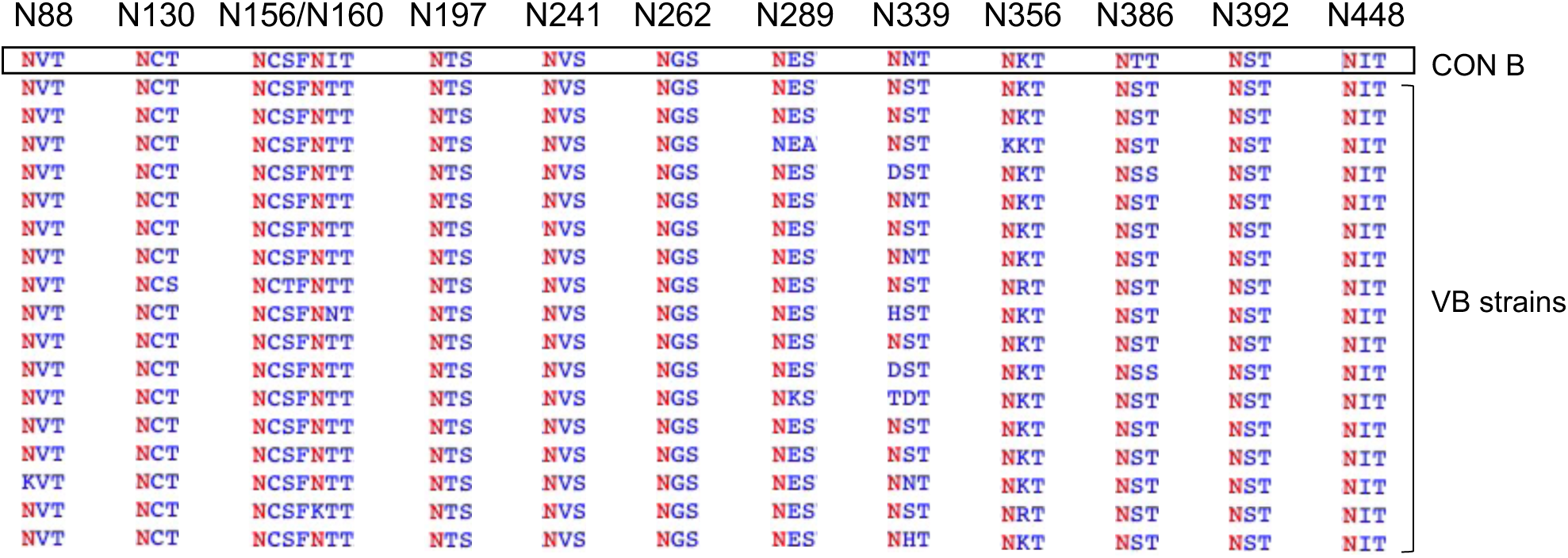
Frequency of N-linked *env* glycan sites in the 17 VB variants. A total of 13 N-linked glycan sites, which are conserved (with >50% frequency) in other subtype B viruses, were analyzed for the 17 VB variants using the N-GlycoSite tool in the Los Alamos HIV Sequence Database. The predicted N-linked glycan sites are highlighted in red. The subtype B consensus sequence (CON B) is used as the reference sequence.

**Figure S3.**
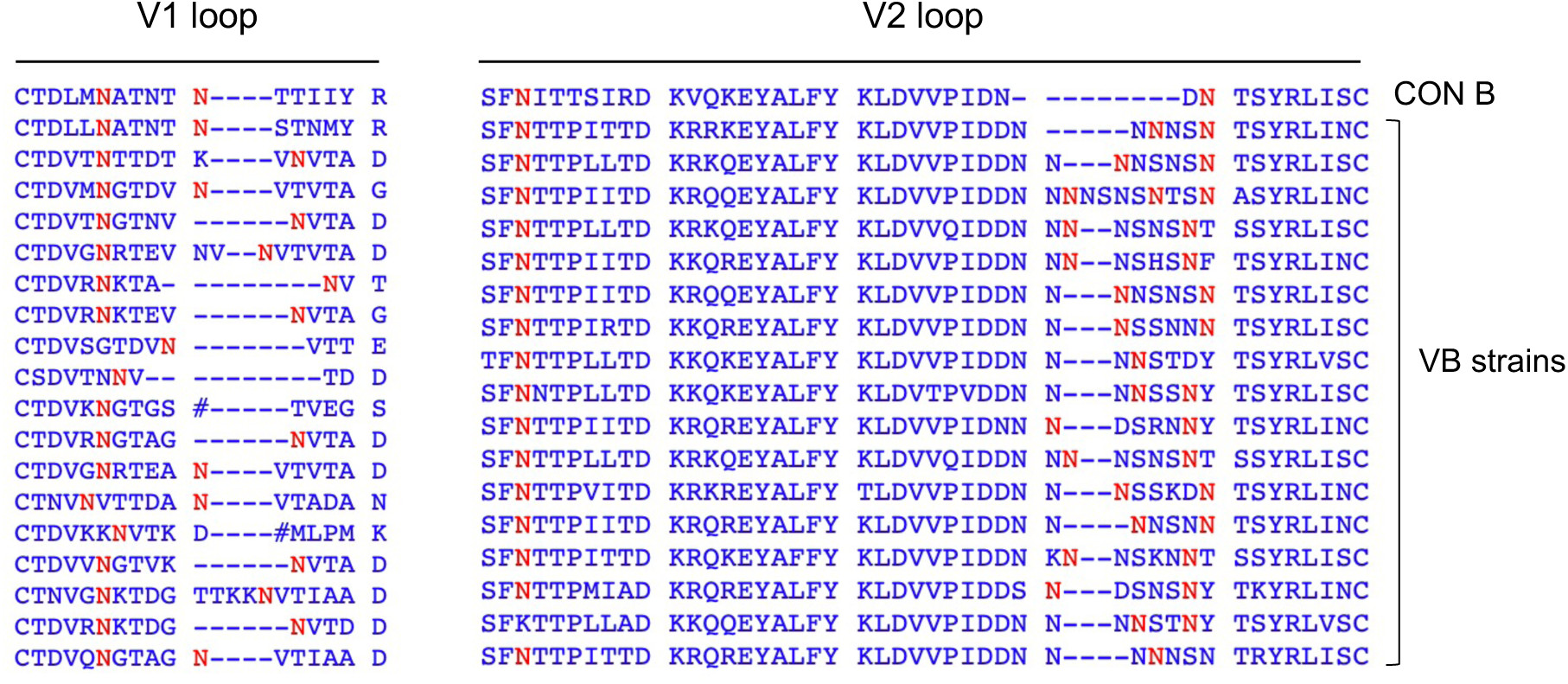
The number of N-linked glycan sites in the V1 and V2 loops of the VB variants as compared to the subtype B consensus sequence. Amino acid alignments of the V1 and V2 loops were obtained by the Gene Cutter tool in the Los Alamos HIV Sequence Database. The N-linked glycan sites in the V1 and V2 loops were analyzed using the N-GlycoSite tool in the Los Alamos HIV Sequence Database. Predicted N-linked glycan sites are shown in red. The subtype B consensus sequence (CON B) is used as the reference sequence.

**Figure S4.**
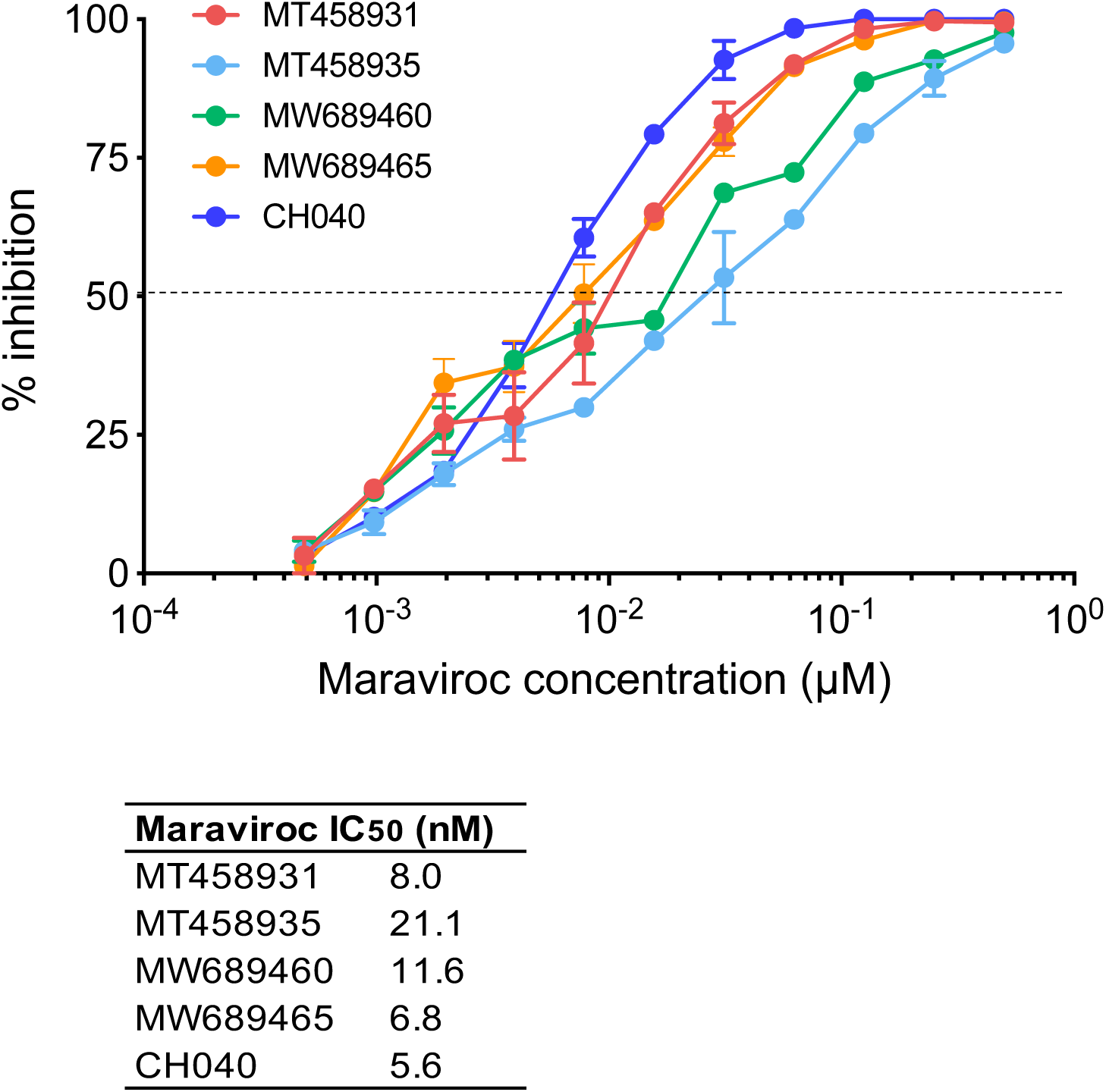
Sensitivity to Maraviroc inhibition in NP-2 CCR5 cell line. Maraviroc sensitivity of four representative VB variants was determined in the NP-2 CCR5 cell line. The subtype B T/F virus CH040 was used as control. The experiments were performed in duplicate, and the error bar represents the standard deviation (SD). The Maraviroc IC_50_ of each virus is shown in the table.

